# Hepatitis C Virus E1 Protein Enhances Macrophage iNOS Expression *In Vitro*

**DOI:** 10.1101/2021.02.15.431273

**Authors:** Batkhishig Munkhjargal, Bilguun Enkhtuvshin, Uranbileg Ulziisaikhan, Baljinnyam Tuvdenjamts, Khulan Unurbuyan, Dolgorsuren Sandagdorj, Tuul Baasandalai, Baasansuren Enkhjargal, Tsogtsaikhan Sandag, Bilegtsaikhan Tsolmon, Enkhsaikhan Lkhagvasuren

## Abstract

**Objective:** Hepatitis C virus (HCV) is a single-stranded RNA virus that causes chronic hepatitis, cirrhosis, and liver cancer. Approximately 170 million individuals are infected with HCV worldwide. The pathogenesis of HCV-associated liver injury is thought to be due to the host antiviral immune response, including the T cell response, and excessive production of proinflammatory cytokines, reactive oxygen species, and nitric oxide (NO).

Interferon-γ (IFN-γ) is a key cytokine in the adaptive immune response that is primarily secreted from CD4+ T helper cells to induce cytotoxic T lymphocyte (CTL) cell response against HCV infection. Another important role of IFN-γ is the activation of macrophages in the liver resulting in inhibition of viral replication and increased NO production.

Enhanced inducible nitric oxide synthase (iNOS) expression and NO production observed in the liver of HCV-infected patients is positively correlated with viral load and hepatic inflammation. HCV-infected macrophages are major producers of NO in the liver. It is not completely understood how HCV proteins affect iNOS expression and what the role of IFN-γ is in HCV protein expression in HCV-infected macrophages. In this study, we examined the effect of INF-γ and HCV proteins on iNOS expression in the Raw264.7 cell line.

**Results:** Consistent with other studies, HCV core and NS5A proteins induced iNOS expression in macrophages. Moreover, HCV E1 protein-enhanced iNOS expression is highest in the presence and absence of IFN-γ activation.

**Conclusion:** These results indicate that hepatitis C virus core, NS5A, E1 protein regulates iNOS protein expression in IFN-γ-activated and resting macrophage cell lines. These findings points to a future research direction for understanding the pathogenesis of HCV-related liver inflammation.

## Introduction

Hepatitis C virus (HCV) is a positive-strand RNA virus belonging to the Flaviviridae family, which has established chronic infection in around 170 million human carriers worldwide [1]. Increasing evidence suggests that nitric oxide (NO) is an important factor in controlling viral infection by affecting the early antiviral immune response in HCV pathogenesis [2]. In the liver of HCV-infected patients, it has been reported that inducible nitric oxide iNOS) expression and excessive local nitric oxide production, which is positively correlated with viral load, produce hepatic inflammation and tissue damage[3].

In many viral infections, iNOS expression appears to be regulated directly by the virus itself or indirectly via interferon-γ (IFN-γ) induction [4,5].

IFN-γ is the primary mediator of HCV-specific antiviral T-cell responses and strongly inhibits HCV replication *in vitro* [6]. IFN-γ is known to upregulate the expression of inducible nitric oxide synthase (iNOS) in monocytes and macrophages, resulting in increased NO production [7]. However, the precise IFN-γ effect on HCV protein expression and crosstalk between HCV proteins and IFN-γ for induction of iNOS expression is not clear.

The primary source of nitric oxide in a liver is activated Kupffer cells (KC) and hepatocytes [8]. It has been shown that HCV proteins regulate iNOS expression and NO production in various cell systems. Many studies have reported that HCV core and non-structural proteins enhance NO production in hepatocytes and endothelial cells. However, there is an inconsistency regarding which viral proteins are responsible because some authors have shown that the core and NS5A proteins are sufficient for iNOS induction in activated KCs. In contrast, Lee et al. found that HCV core protein inhibits NO production in cultured Raw264.7 and J774 cell lines [2,9].

To understand these differences, in this study we examined the effects of HCV proteins on iNOS expression in IFN-γ-activated and resting-state macrophages.

## Materials and Methods

### Cell culture

The murine macrophage cell line RAW264.7 (Core laboratory, MNUMS, Ulaanbaatar Mongolia) was used in this study. RAW264.7 cells were cultured in RPMI 1640 (Cat: 12-702F from Lonza Pharma & Biotech, USA) supplemented with 10% inactivated fetal calf serum (FCS) (Cat: a3520501 Thermo Fisher Scientific Inc, USA), an 1% antibiotic mixture (Cat: 15140-122 from Thermo Fisher Scientific Inc, USA). Cells were incubated at 37°C and humidified in 5% CO2 until cell growth reached 85% of the culture plates’ surface area.

### Transfection and IFN-γ treatment

The cells were seeded in a 6-well plate and were cultured until they reached a confluence of 70%. The culture was starved of nutrients by with holding the serum overnight before transfection. Transfection of 1 µg pHCV-Core, pHCV-NS5A and pHCV-E1 (Cat: VG40278-UT, VG40284-UT, VG40279-UT from Sino biological, HK) plasmids were performed in separate wells using Lipofectamine 3000 (Cat: L3000001 from Invitrogen, USA) and cultured in a complete medium for 48 hours. Half of the cultures were then treated with IFN-γ (25 ng/ml) (Cat: 315-05-100 from PeproTech Rocky Hill, USA) for 18 hours at 48 hours post-transfection[10].

### Immunoblotting

The cells were lysed using RIPA buffer supplemented with protease and phosphatase inhibitors (Lot: 06131601 from Thermo Fisher Scientific Inc, USA). The protein concentration was measured by Pierce BCA protein assay kit (Cat: 23228 from Thermo Fisher Scientific Inc, USA) according to the manufacturer’s instruction. Proteins were resolved by SDS electrophoresis and transferred onto nitrocellulose membranes. The membrane was incubated for 24 hours with the HCV anti-core, anti-E1, anti-NS5A (Cat: ab18929, ab13833, ab54555 from Abcam, UK) anti-iNOS and anti-p38 antibodies (Cat: 2982S, 9215S from Cell Signaling Technology, USA). The blot was then incubated with the corresponding HRP-conjugated secondary antibodies (Cell Signaling Technology, USA Cat: 111-035-003). The proteins were visualized using the ECL system (Lot OA183335 from Pierce, Rockford, USA). Total p38 expression was used as an internal control [11].

### Ethical statement

The study was approved by The Ministry of Health (MoH)-Medical Ethics Committee (№80).

## Results

### Effect of IFN-γ on the HCV proteins expression in macrophage

Kupffer cells (KCs) are liver macrophages and represent 15 to 20% of the total liver cell population. Classical type macrophage activation against HCV occurs by T cell-mediated immunity in the presence of IFN-γ. It has been shown that IFN-γ suppresses HCV replication through activation of various ISG genes. However, IFN-γ’s direct impact on HCV protein expression is not well studied. We examined the effect of IFN-γ on HCV proteins using transient transfection of HCV core, NS5A and E1 protein-coding plasmids. The experimental procedure for testing the effect of IFN -γ on HCV protein expression is as outlined in (Figure 1A). Raw264.7 cells were seeded 2 days before the transient transfection with equal amounts of HCV-core, NS5A and E1 coding plasmids driven by a CMV promoter. HCV E1, NS5A and core proteins are detected around 31 kD, 56 kD and 21 kD, respectively, by Western blot (Figure 1B). Our Western blot results proved (Figure 1B) that we successfully transfected DNA to RAW264.7 cells. IFN-γ treatment was performed at 2 days post-transfection to test the HCV protein expression in response to IFN-γ-mediated macrophage activation. An increase in HCV E1 protein expression was observed when the culture medium was supplemented with IFN-γ. In contrast, HCV core and NS5A expressions were inhibited by IFN-γ treatment (Figure 1B).

**Figure 1.**
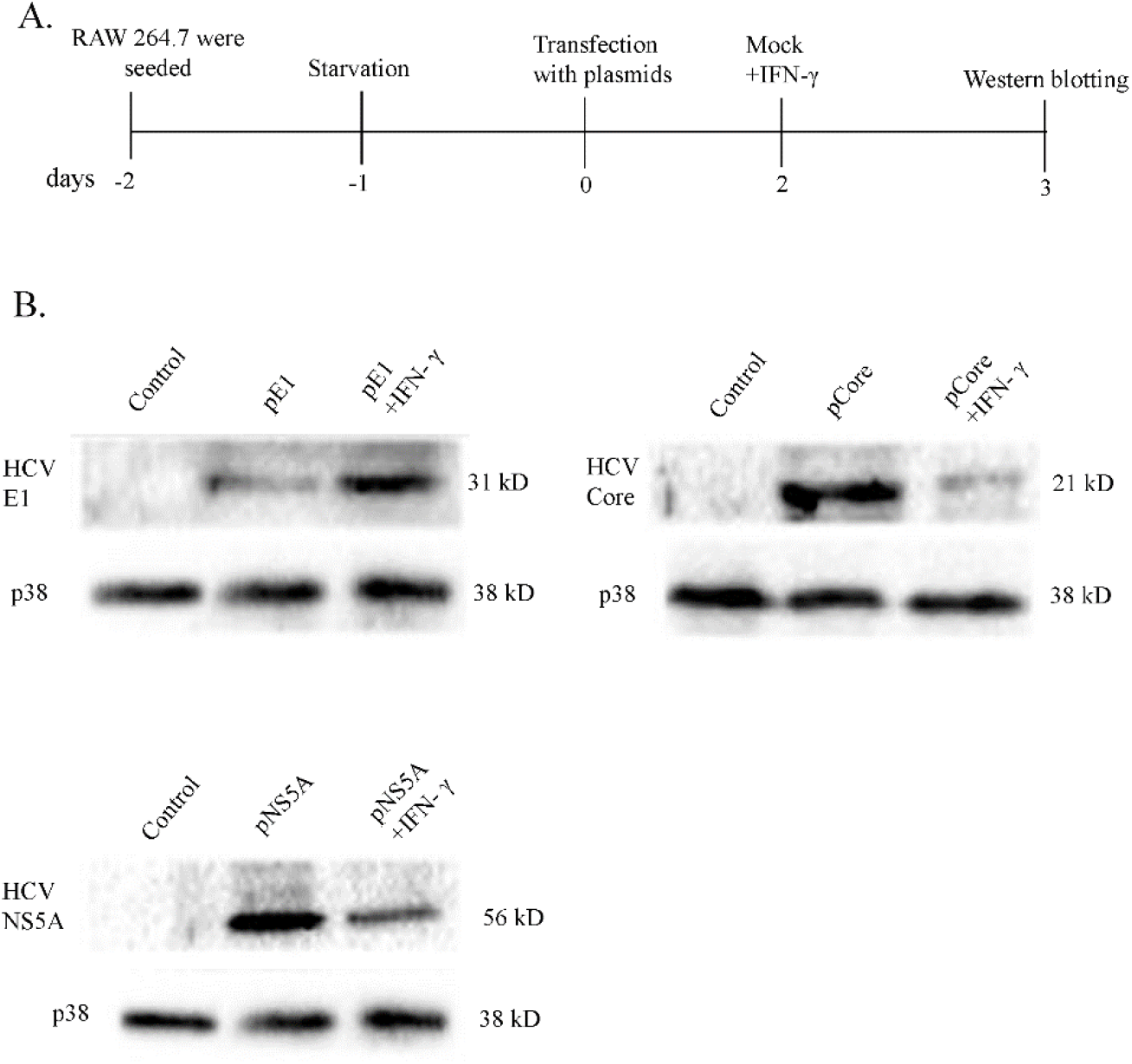
Effect of IFNγ on the HCV proteins expression in macrophages. (A) Illustration showing the experimental design. (B) Non-transfected RAW264.7 cells served as a negative control (Lane 1). The expression of HCV Core, NS5A, E1 protein in RAW264.7 cells was detected by Western Blot which indicates successful transfection (Lane 2). HCV core, NS5A, and E1 protein expressions were examined after IFN-γ treatment (Lane 3). Total p38 expression served as a loading control.

### HCV E1 protein modulates iNOS expression in Raw264.7 cells

It is well known that HCV infection upregulates iNOS expression in *vitro* and in *vivo*, albeit the responsible HCV protein for increased iNOS expression in macrophages remains elusive. The increased iNOS expression and NO production induce liver tissue damage and have no impact on HCV replication and viral protein expression. Next, we asked how IFN-γ mediated the HCV-induced upregulation of iNOS by the host immune response. The experimental procedure is outlined (Figure 2A.). The RAW264.7 cells were starved of serum 24 hours before transfection to synchronize all cells into the same phase of the cell cycle and to eliminate possible noises of iNOS expression. IFN-γ and mock treatments were done at 2 days post-transfection with viral protein-coding plasmids. The expression of iNOS protein was detected by Western Blot using an anti-iNOS antibody at day 3 post-transfection. Mock-transfected resting RAW264.7cells showed no detectable iNOS expression by western blot (Lane 1). IFN-γ treatment enhanced iNOS expression slightly (compare Lane 1 and Lane 2). HCV core transfection resulted in marginally increased expression of iNOS while no detectable signal was observed in the NS5A transfection (compare Lane 3 and 7 to Lane 1). To our surprise, HCV E1 transfection increased iNOS expression (compare Lane 1 and 5). The combination of HCV proteins transfection and IFN-γ mediated macrophage activation induced more iNOS expression compared to IFN-γ treatment alone (compare Lane 4, 6 and 8 to Lane 2). iNOS expression was markedly increased when with HCV E1 plasmid transfection was combined with IFN-γ treatment (compare Lanes 5 and 6) (Figure 2B).

**Figure 2.**
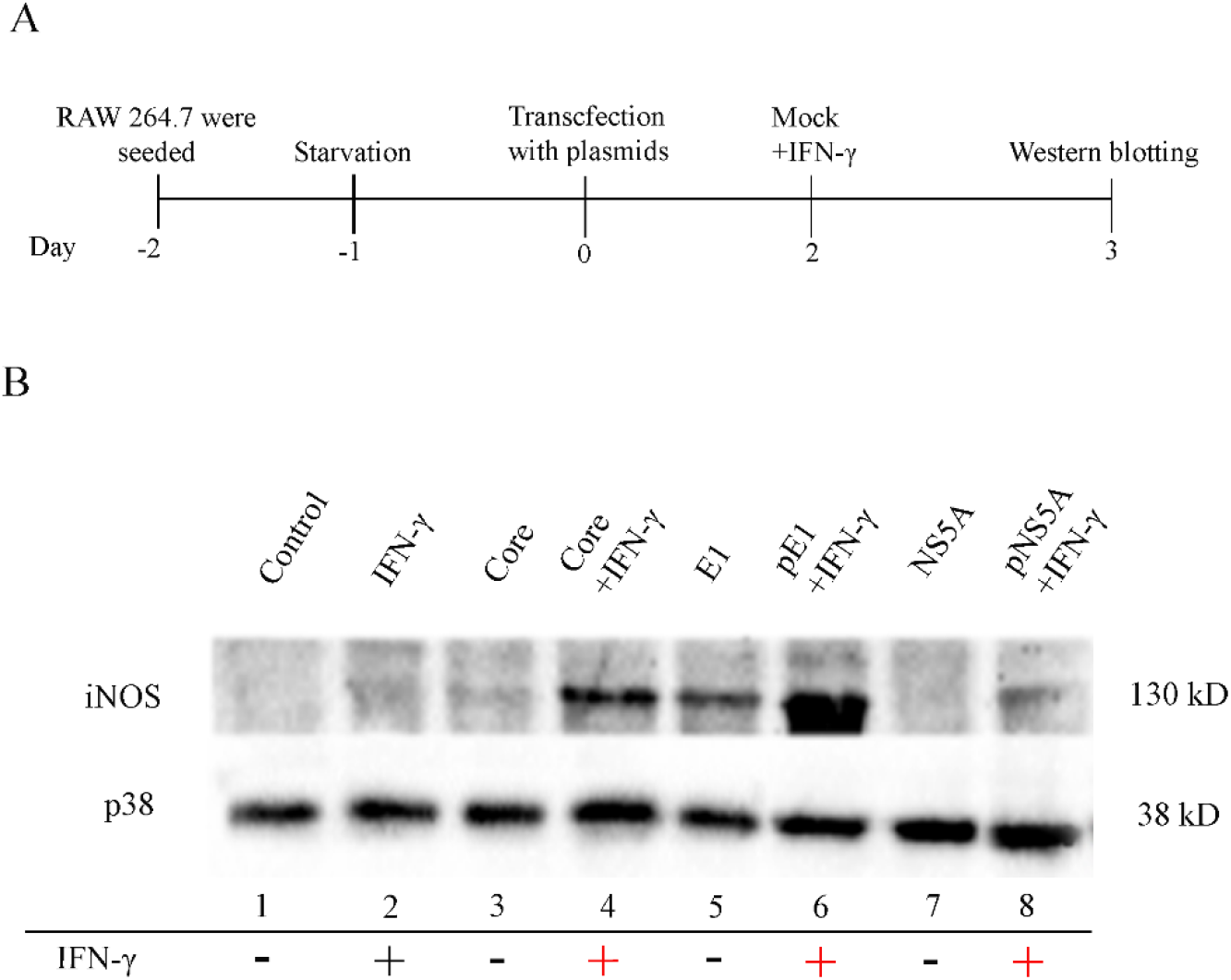
HCV proteins modulate iNOS expression in Raw264.7 cells. Illustration showing the experimental design. (B) iNOS expression in Raw264.7 cells was detected by Western blot analysis using anti-iNOS antibodies (mouse). Non-transfected RAW264.7 cells served as a negative control (Lane 1). Non-transfected cells were activated by 25ng/ml IFN-γ (Lane 2). iNOS expression in RAW264.7 cells was detected by Western Blot after transfection with pHCV-Core, pHCV-E1 and pHCV-NS5A plasmids without IFN-γ activation (Lanes 3,5,7). HCV core, E1, NS5A protein-expressing RAW264.7 cells were activated by 25ng/ml IFN-γ (Lanes 4,6,8). Total p38 expression served as a loading control. (+) treated with IFN-γ, (-) untreated with IFN-γ

## Discussion

In this study, we revisited the relationship between IFN-γ, HCV proteins and iNOS expression in macrophages using a transient transfection system. Our Western blot results proved successful transfection of HCV E1, NS5A and core protein-coding plasmids (Figure 1B). IFN-γ treatment on HCV core and NS5A protein-expressing RAW264.7cells showed inhibition of viral protein expression. In contrast, HCV E1 protein expression was increased when cells were treated with IFN-γ (Figure 1B).

iNOS expression were influenced positively increases transfection of HCV proteins with the IFN-γ treatment. Surprisingly, we found that HCV E1 protein upregulates iNOS expression *in vitro* most strongly (Figure 2B).

The HCV protein’s effect on NO production and iNOS expression has been studied in various cell systems in earlier reports (Table 1). HCV core protein increased iNOS expression in the studies using CHL, Raji, HepG2, primary human conjunctival fibroblast, human corneal epithelial cell lines and mouse liver [12-17]. In contrast, iNOS expression was downregulated by HCV core protein in the in vitro studies using Raw264.7 and J774 cell lines and coculture system of LSECs+HepG2 [18,19]. The opposite effect of HCV core protein on iNOS expression between this study and earlier reports can be related to several possibilities: 1)We detected increased iNOS expression when HCV core protein was treated together with IFN-γ in Raw264.7 cells, while earlier reports observed decreased NO production when co-treated with HCV core, LPS and Zymosan in Raw264.7 and J774 cells. 2) Earlier reports used the coculture system of LSECs+HepG2 to study the effect of HCV core protein on iNOS expression. Using a different cell culture system could be a cause of opposite observations. iNOS expression was upregulated by HCV NS5A protein in HepG2 and CHL cell systems [12,20].

**Table 1.** Previous reports of HCV protein’s effect on NO production and iNOS expression.

| Author | Experimental model | HCV protein | iNOS expression | NO production |
| --- | --- | --- | --- | --- |
| Keigo Machida et al. [17] | Mouse liver | Core | Upregulated |  |
| Maria Victoria Garcia et al. [12] | CHL cell line | Core<br>NS5A | Upregulated<br>Upregulated |  |
| Keigo Machida et al. [13] | Raji cell | Core | Upregulated |  |
| Susana de Lucas et al.[15] | HepG2 | Core | Upregulated |  |
| Chu Hee Lee et al.[18] | Raw 264.7 and J774 | Core |  | Downregulated |
| Li-Jie Sun et al.[19] | LSECs+HepG2 | Core |  | Downregulated |
| Ayilam Ramachandran Rajalakshmy et al. [15] | Primary human conjunctival fibroblasts | Core | Upregulated | Upregulated |
| Yong-Fang Jiang et al.[20] | HepG2 | NS5A | Upregulated |  |
| Ayilam Ramachandran Rajalakshmy et al. [16] | Human corneal epithelial cell line | Core | Upregulated | Upregulated |

We also found that HCV NS5A could stimulate iNOS expression in the Raw264.7 cell line. We were unable to find any studies about HCV E1 influence on iNOS expression in macrophages in our literature review (Table 1). Approximately up to 85% of acute HCV infections result in persistent infection and can lead to life-threatening chronic conditions, including cirrhosis and hepatocellular carcinoma [21]. The mechanism of chronicity is explained by an insufficient HCV-specific CTL response, suppressed type 1 helper T cell response and generation of viral escape mutations [22,23].

Frese et al. demonstrated HCV viral protein synthesis and replication were not suppressed by increased iNOS expression and NO in hepatocytes. They also reported that in iNOS-deficient mice, increased production of NO might weaken T cell antiviral responses by impairing T helper 1 cell activity, leading the virus to overcome the pressure of the T cell immunity [24]. Viral escape mutations are frequently another important reason for viral persistence in HCV infection, and evidence suggests that NO could play a role in this scenario by accelerating the mutation rate of HCV RNA during viral infection in vivo [23]. What could then be the HCV E1 protein’s role in the pathogenesis of HCV persistence? Is it possible that HCV E1 protein, driven iNOS expression, may play a role in HCV persistence? HCV E1 protein expression is increased when IFN-γ (T helper I cytokine) is added (Figure 3). HCV E1 protein enhances iNOS expression in IFN-γ activated macrophages. Enhanced iNOS expression resulted in excessive NO production in Kupffer cells. It may be that the HCV E1 protein indirectly modulates T helper 1 cell activity and accelerates HCV mutation rate (Figure 3). The HCV E1 protein allows HCV to escape and weaken B and T cell immune response.

**Figure 3.**
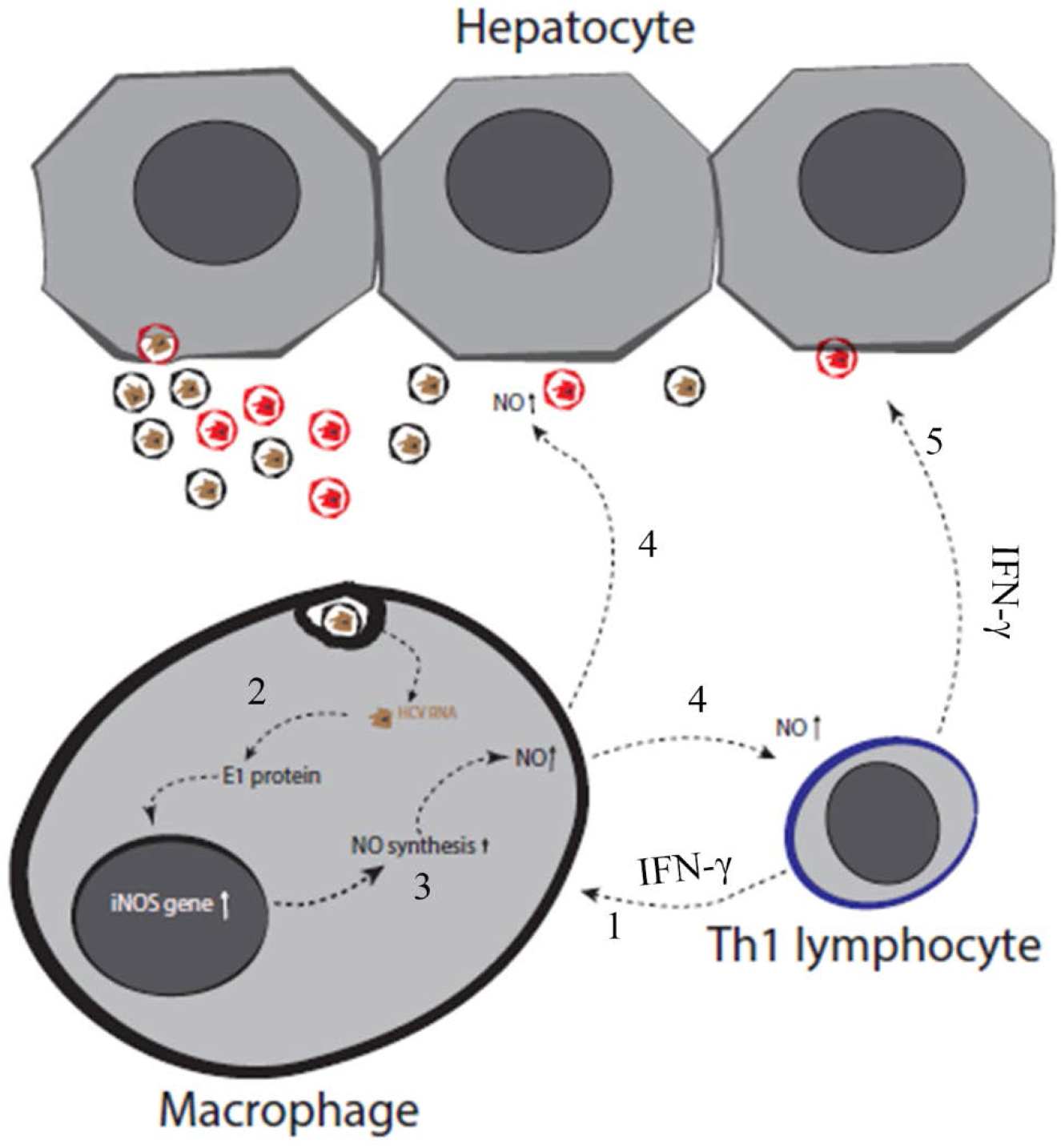
Possible mechanism of HCV E1 protein role on HCV persistence. HCV, viral proteins, liver cells and signalling molecules involved in the pathogenesis of HCV persistence are illustrated. HCV persistence mechanism is generally explained by suppressed T helper 1 response and viral escape mutations. Nitric oxide induces weakens T helper 1 response and accelerates virus mutations (illustrated by block arrow, step 4, 5) (references). IFN-γ secreted from HCV specific T helper 1 lymphocyte induces HCV E1 protein expression in HCV-infected Kupffer cells (step 1). HCV E1 enhances iNOS expression in Kupffer cells (step 2). IFN-γ activated HCV E1 protein in Kupffer cells results in a further increase of iNOS expression (step 3). Excessive nitric oxide produced from HCV E1-derived iNOS activation weakens the T helper 1 response (step 4) and leads to the generation of escape mutations (step 5).

In summary, HCV core and NS5A proteins induce iNOS expression when combined together with IFN-γ. IFN-γ alone inhibited the expression of those viral proteins. However, HCV E1 protein strongly induced iNOS expression with or without IFN-γ. Surprisingly, IFN-γ treatment further elevated HCV E1 protein levels. This is the first report to our knowledge that HCV E1 protein drives enhanced expression of iNOS in the macrophage cell line. Our results clarify there is crosstalk between HCV proteins and iNOS expression in the background of IFN-γ treatment. HCV E1-induced iNOS expression may modulate HC-associated tissue damage pathogenesis and viral persistence. These findings update our understanding of HCV pathogenesis and HCV persistence.

## Acknowledgments

The research funding was provided by the Mongolian Foundation for Science and Technology and the Ministry of Education, Culture, Science and Sports of Mongolia (Basic research grant: 2018/39). We thank Lkhagvasuren Damdindorj for technical support. We are grateful to Dr. Ron Anderson for the critical reading of the manuscript.

